# Non-Invasive Diagnostic Evaluation of Urinary Exosomal Let-7c Cluster Expression in Bladder Cancer Using Machine Learning Approaches

**DOI:** 10.1101/2025.09.18.677033

**Authors:** Sukhad Kural, Lalit Kumar, Abhay Kumar Pathak, Shweta Singh, Garima Jain, Mahima Yadav, Yashasvi Singh, Ujwal Kumar, Sameer Trivedi, Manjari Gupta, Vibhav Gautam, Parimal Das

## Abstract

**Background:** Bladder cancer (BCa) diagnosis typically relies on invasive cystoscopy, which is effective but costly and uncomfortable. Urinary microRNAs (miRNAs), especially exosomal ones, are promising non-invasive biomarkers due to their stability in biological fluids and disease specificity. However, challenges such as population variability, methodological inconsistencies and normalization issues hinder their clinical translation, emphasizing the need for innovative approaches to enhance diagnostic performance.

**Objective:** To evaluate the diagnostic potential of urinary exosomal let-7c cluster (let-7c-5p, miR-99a-5p and miR-125b-5p) in BCa patients by integrating miRNA expression data with Machine Learning (ML) models.

**Methods:** Urine samples were collected from 66 participants, including 50 BCa patients and 16 healthy controls (HC). Exosomal miRNAs were isolated and quantified using Quantitative Real-Time-Polymerase-Chain-Reaction (qRT-PCR). Statistical analysis and hypothesis tests were conducted to explore the nature and diagnostic relevance of individual biomarkers. A logistic regression classifier was applied to evaluate both the combined and differential diagnostic capabilities of the selected biomarkers. Accuracy, precision, recall and AU-ROC scores were used to assess model performance. Bioinformatics analysis was performed to identify pathways associated with the features prioritized by the ML models, ensuring their relevance to BCa.

**Results:** The result revealed significant differentiation between BCa patients and HC, with miR-99a-5p (p=0.013, AU-ROC=0.71) and miR-125b-5p (p=0.047, AU-ROC=0.64) demonstrating reliable diagnostic performance (let-7c-5p showed weaker discrimination, AU-ROC=0.65, p>0.1). The logistic regression ML model achieved an accuracy of 80.0% (AU-ROC=0.86, recall=100%) in distinguishing cancer from HC and 53.3% (AU-ROC=0.63) when applied to miRNA-only grade classification. When clinical variables were integrated with miRNA expression, performance improved to 73.3% accuracy (AU-ROC=0.61) for high-versus low-grade differentiation. Across Ta–T2, miR-99a-5p displayed relatively better separation, whereas let-7c-5p and miR-125b-5p showed weak stage-related differences. The integration of bioinformatics analysis confirmed the biological relevance of these miRNAs in BCa-related pathways, including PI3K–Akt, p53, NF-κB and RAS/MAPK signaling, with hub genes such as TP53, MYC, EGFR, and CCND1 identified, further validating the diagnostic utility of the selected biomarkers.

**Conclusion:** Urinary let-7c cluster miRNAs demonstrate promising diagnostic potential when analyzed with ML models, offering a non-invasive alternative to conventional methods. These findings highlight the promise of ML-based approaches alongside molecular markers for advancing clinical diagnostics in BCa.

## 1. Introduction

Bladder cancer (BCa) accounts for around 3–4% of cancers in men, ranking as the fourth leading cancer in males and the ninth most common globally. Despite advances in management, it is characterized by high recurrence rates and poor prognosis in advanced stages^1^. The risk factors are diverse and include tobacco use, exposure to dyes and chemicals, chronic urinary tract infections (UTIs) and long-term bladder irritation caused by the presence of catheters or bladder stones^2,3^. Few other factors include aging, gender, genetic predisposition and conditions like chronic cystitis^4^. Painless gross hematuria is the most common symptom seen in 80-90% of patients, but only 10% of those with hematuria are diagnosed with BCa^5^. BCa is a heterogeneous disease, with more than 90% of cases arising as urothelial carcinoma. Less common histological variants include squamous cell carcinoma, adenocarcinoma and neuroendocrine tumors^6^. Based on the extent of muscle invasion, BCa is categorized as non–muscle invasive BCa (NMIBCa; stages Ta, T1, CIS) or muscle-invasive BCa (MIBCa; stage T2 and above)^7^. Approximately 75% of cases are non-muscle invasive and are primarily managed with transurethral resection of bladder tumor (TURBT), chemotherapy and intravesical Bacillus Calmette-Guérin (BCG) therapy that contribute to favorable therapeutic results^8,9^. However, MIBC is typically treated with a combination of radical cystectomy (removal of the bladder) and systemic chemotherapy (including cisplatin-based regimens), sometimes followed by adjuvant therapy^10^. Although cystoscopy and urinary cytology remain the gold standards for BCa diagnosis, the invasive nature of cystoscopy and the limited sensitivity of cytology in low-grade (LG) tumors pose significant challenges^11^. As a result, both methods often fail to detect malignancies at an early stage, when treatment is most effective. Moreover, these approaches provide little information about the molecular characteristics of tumors, which are increasingly relevant for guiding therapy, underscoring the need for more effective non-invasive alternatives.

Extracellular vesicles (EVs) are a group of membrane-bound vesicles secreted by all eukaryotic cells and found in a variety of biofluids, including urine, functioning as molecular cargo carrying diverse biomolecules such as mRNAs, microRNAs (miRNAs), lncRNAs and proteins, making them great source of biomarkers^12^. Exosomes are a subtype of EVs between 30 and 150 nm in size derived from endosomes that transfer biomolecules to facilitate cell communication and play a role in cancer progression and metastasis^13^. miRNAs are remarkably stable in adverse conditions, such as acidic or alkaline pH, thermal stress and RNase activity^14^. When enclosed in urinary exosomes, they offer a non-invasive and accessible approach for detecting and monitoring BCa, with strong clinical potential. Approximately 99.96% of urinary exosomes are released by cells from the kidney, urinary tract lining and male urogenital tract^15^. This direct exosome shedding by urological tumors into the urine makes them highly promising candidates for sensitive and urological cancer biomarkers^16^.

Liquid biopsy has emerged as an effective tool for early screening, detection of tumors and for monitoring disease progression. Over the last decade, miRNAs have emerged as promising candidate for liquid biopsy based diagnostics due to their stability in biofluids, tissue specificity and key regulatory roles in cancer-associated pathways^17^. miRNAs are short, ∼22-nucleotide, noncoding RNAs that modulate gene expression after transcription by binding to the 3′ untranslated regions (3′ UTRs) of target mRNAs, leading to repression of translation or mRNA degradation^18^. Gene expression is largely influenced by miRNAs and their imbalance is frequently observed in cancer^19^. Some miRNAs promote tumor development when overexpressed by suppressing genes that inhibit growth (e.g., miR-21, which targets PTEN and PDCD4 to enhance cell proliferation and survival in urologic cancers, including BCa^20^; miR-155, which inhibits DMTF1, contributing to BCa progression^21^), while others that act as tumor suppressors are often downregulated and allow cancer cells to proliferate unchecked (e.g., the let-7 family, whose reduction leads to RAS overexpression and promotes tumorigenesis in BCa^22^).

The let-7 family of miRNAs is known for its tumor-suppressive role and is significantly involved in regulating the cell cycle, differentiation and apoptosis^23^. As a tumor suppressor miRNA, let-7c is well known to regulate oncogenes such as HMGA2^24^, H-Ras^25^ and c-MYC^26^, thereby contributing to suppression of tumor progression^27^. The let-7c/miR-99a/miR-125b cluster, located on chromosome 21, is a well-conserved group of miRNAs known for their tumor-suppressive functions across a range of cancers^28^, including BCa, hepatocellular carcinoma, prostate cancer, cholangiocarcinoma, mesothelioma and other malignancies. These miRNAs collectively target multiple cancer pathways such as mTOR, IGF1R and STAT3 modulating key processes like proliferation, apoptosis and cellular stemness^28–30^. In BCa, these miRNAs are typically downregulated in tumor tissues, which is consistent with their role in inhibiting cancer progression^31,32^. Given their ability to regulate multiple oncogenic pathways and the strength of existing evidence, the let-7c cluster was selected for analysis in BCa as a potential non-invasive biomarker for diagnosis and monitoring disease progression.

Machine Learning (ML), a branch of Artificial Intelligence (AI), enables systems to learn from data without being explicitly programmed. Instead of relying on explicit coding, ML models leverage algorithms to uncover patterns in data, which enables predictive and decision-making capabilities. Supervised, unsupervised, semi-supervised, and reinforcement learning form the four categories of ML models, each fulfilling different functions^33^. As these models process more data, they improve over time, making ML a powerful tool across many domains, including healthcare^34,35^.

In biological research, ML is revolutionizing the way we analyze complex datasets such as genomics, proteomics and medical imaging^36^. Deep biological datasets like sequencing are often large, noisy, and nonlinear, which is where ML shines. For example, ML has been widely used in cancer diagnostics, particularly in biomarker discovery, by analyzing RNA sequencing or microarray data^37^. However, despite its successes, supervised learning models especially in analyzing Quantitative Real-Time Polymerase Chain Reaction (qRT-PCR) data are still underutilized. Few studies have effectively leveraged qRT-PCR data with supervised learning to improve diagnostic accuracy^38^. This study presents one of very few studies with integrative demonstration of supervised ML applied directly to qRT-PCR derived profiling in cancer, highlighting a clinically scalable and cost-effective approach that advances the translational potential of liquid biopsy diagnostics.

Linear regression, which assumes a straight forward relationship between inputs and outputs, is frequently used but has its limitations^39^. Logistic regression, in contrast, is a widely used supervised model that performs effectively on binary classification tasks, such as identifying diseased versus healthy conditions^40^. This algorithm estimates the probability of a binary outcome using a logistic function, making it ideal for analyzing diagnostic data like qRT-PCR results^41^. While logistic regression is simple, it can be very powerful when combined with proper feature selection and normalization. Additionally, its interpretability makes it valuable in biological research, where understanding the significance of each feature is essential.

To strengthen these findings, we conducted bioinformatics analyses including target prediction, overlap with BCa-specific genes, functional enrichment and PPI network construction. These analyses demonstrated that the dysregulated miRNAs identified in our cohort are linked to established pathways of BCa progression, thereby reinforcing their biological relevance. By integrating experimental data with in silico validation, our study provides supportive evidence for the development of a non-invasive, exosome-based diagnostic framework for BCa.

### 1.1. Research Gap

Even though research on BCa has advanced significantly, non-invasive diagnostic techniques have not been widely adopted as alternatives to invasive procedures in clinical settings and their current application is still primarily complementary rather than independent. For example, a number of urine-based tests that have received U.S. Food and Drug Administration (FDA) approval, including Nuclear Matrix Protein 22 (NMP22) BladderChek, Bladder Tumor Antigen (BTA) stat and UroVysion Fluorescence In Situ Hybridization (FISH), which proves potential and clinical relevance of non-invasive testing^42^. However, these tests have drawbacks, such as NMP22 and BTA stat shows high false positives in the presence of hematuria (blood in urine) or inflammation, while UroVysion FISH, though more effective in HG disease, performs poorly in LG cases and is also costly and technically demanding, which has limited their use to supportive tests rather than replacements for cystoscopy, with urine cytology often used as a complementary test^43^. Although there are few studies which investigated the let-7c cluster in BCa^32,44^, not enough is known about its diagnostic aspect in urinary exosomes. Moreover, there has been limited study of qRT-PCR derived miRNA expression data examined using supervised ML models in the current literature, which has concentrated chiefly on high-throughput platforms like microarrays and RNA sequencing for biomarker discovery^38,45^. In this study, we explored the potential of applying supervised ML to qRT-PCR derived urinary exosomal miRNA data, an approach being more practical and accessible than sequencing-based platforms. Incorporating ML offers the ability to capture subtle fluctuations in expression features that conventional linear statistical methods may overlook, thereby enhancing the diagnostic value of this widely used technique.

### 1.2. Research Questions

This study examines the urinary exosomal miRNAs, particularly the let-7c cluster as non-invasive biomarkers for BCa by combining molecular diagnostics, bioinformatics and ML. The following research questions guided the investigation:

- Assess whether let-7c cluster members (let-7c-5p, miR-99a-5p, miR-125b-5p) are differentially expressed in BCa patients compared with healthy controls (HC), as well as across different tumor grades and stages.
- Determine the diagnostic performance of qRT-PCR derived miRNA expression data using supervised ML models.
- Explore associations between miRNA expression and clinical or lifestyle variables such as tumor grade, stage, size, and tobacco use.
- Elucidate the biological pathways and molecular targets linked to these miRNAs through integrative bioinformatics analyses (PPI networks, GO/KEGG enrichment).

The purpose of raising such questions is to integrate classical biomarker discovery methods with advanced ML approaches through the development of an affordable and accessible diagnostic model for BCa. By utilizing exosomal miRNA expression, this study aims to facilitate the development of precision-based, non-invasive diagnostics with clinical relevance.

## 2. Methods

### 2.1. Study design

This prospective cohort study, involving 50 patients with BCa and 16 HC, was performed at the Department of Urology in association with the Departments of Radiology, Pathology, and the Centre for Genetic Disorders. The study was conducted following approval from the Institutional Ethics Committee (EC/3411), with written informed consent obtained from all enrolled participants. A total of 50 patients (≥18 years old) suspected of having bladder tumors who underwent primary TURBT were enrolled between October 2022 and April 2024. 16 healthy, age and sex-matched control subjects with no known urological pathology were recruited for comparative analysis of urinary let-7c cluster expression. Exclusion criteria included tumors arising in bladder diverticula, incomplete medical documentation, history of TURBT, prior chemoradiotherapy, impaired renal function and earlier pelvic malignancy. Patients who did not consent for the study were also excluded.

### 2.2. Sample Collection and Exosomal RNA Isolation

Approximately 25–30 mL of midstream early-morning urine was collected pre-operatively from patients undergoing TURBT using sterile urine collection vials. Samples were transferred to the laboratory within two hours of collection in a temperature-maintained (4 °C) insulated box, without any intermediate storage or freezing. A volume of 10 mL from each sample was utilized for exosomal RNA extraction using the Norgen Urine Exosome RNA Isolation Kit (Cat#47200, Norgen Biotek Corporation), following the manufacturer’s protocol. A total of 20 µL of Elution Solution A (kit-provided) was used to elute RNA, whose concentration and purity were measured with a NanoDrop OneC UV-Vis Spectrophotometer (Thermo Fisher Scientific, USA) before being stored at −80 °C for downstream analyses.

### 2.3. Complementary DNA (cDNA) synthesis and qRT-PCR for Exosomal miRNA

Using the RevertAid First Strand cDNA Synthesis Kit (Thermo Scientific, Catalog #K1622, USA), complementary DNA (cDNA) synthesis of exosomal RNA was performed. Custom stem-loop primers were introduced to the reverse transcription reaction in order to attain specificity against miRNA targets. **Table 1** summarizes the full set of primer sequences utilized in the study. qRT-PCR experiments were performed on the QuantStudio 6 Flex Real-Time PCR System (Applied Biosystems/Thermo Scientific, USA). 1 μL of diluted 1:5 cDNA, 6.25 μL of Maxima SYBR Green/ROX qPCR Master Mix (2X; Catalog #K0221, Thermo Scientific, USA), 1 μL of each forward and reverse primer (10 μM) and nuclease-free water for the final volume were all included in each 12.5 μL reaction. The thermal cycling technique included a 10 minute denaturation stage at 95°C, followed by 40 amplification cycles of 15 seconds at 95°C and 1 minute at 60°C. The overall experimental workflow is illustrated in **Figure 1A**.

**Figure 1:**
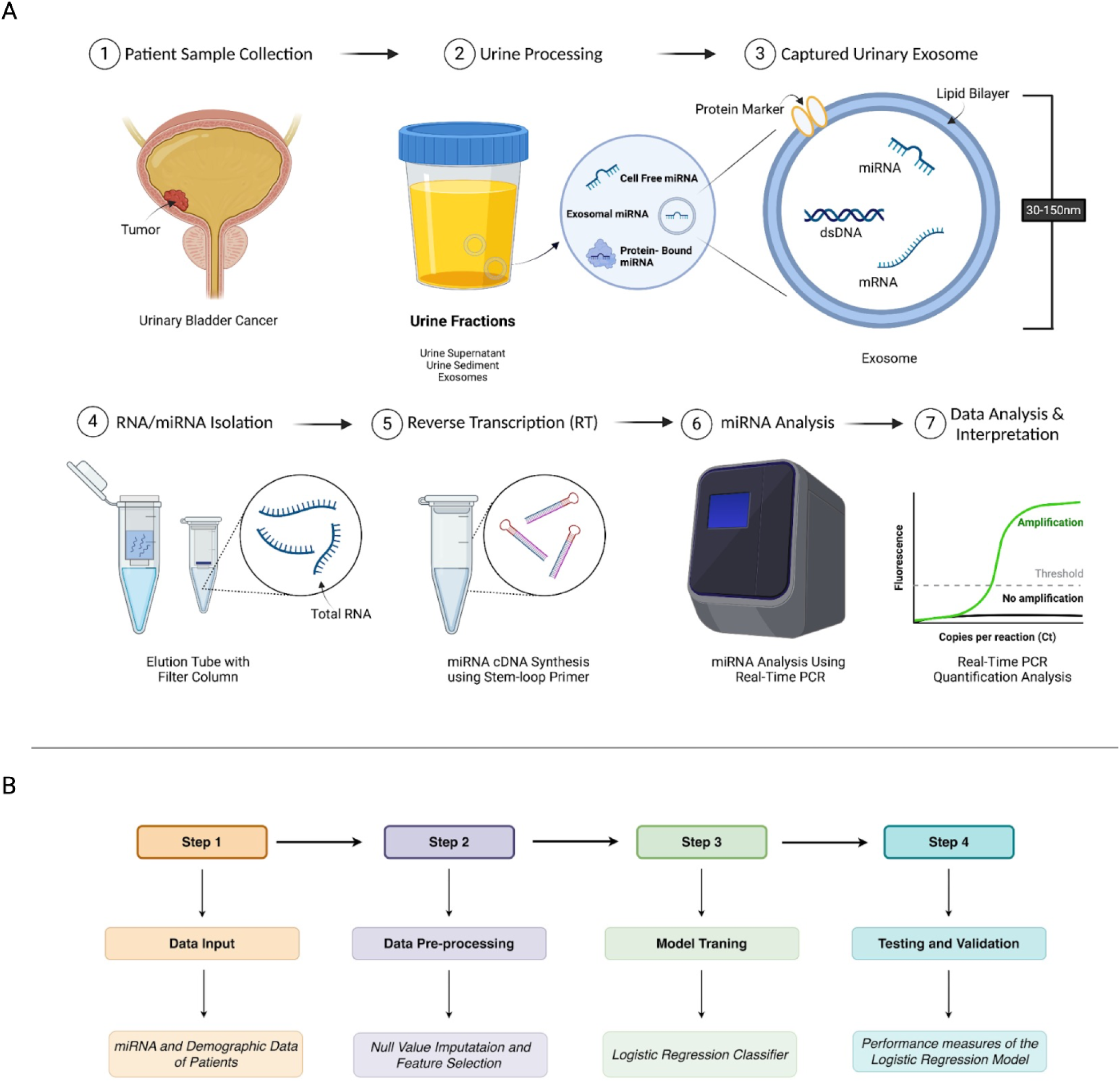
**A.** Experimental workflow illustrating the methodology used for urinary exosomal miRNA profiling in BCa. **B.** Flowchart of the model developed (Various Classification ML)

**Table 1:**
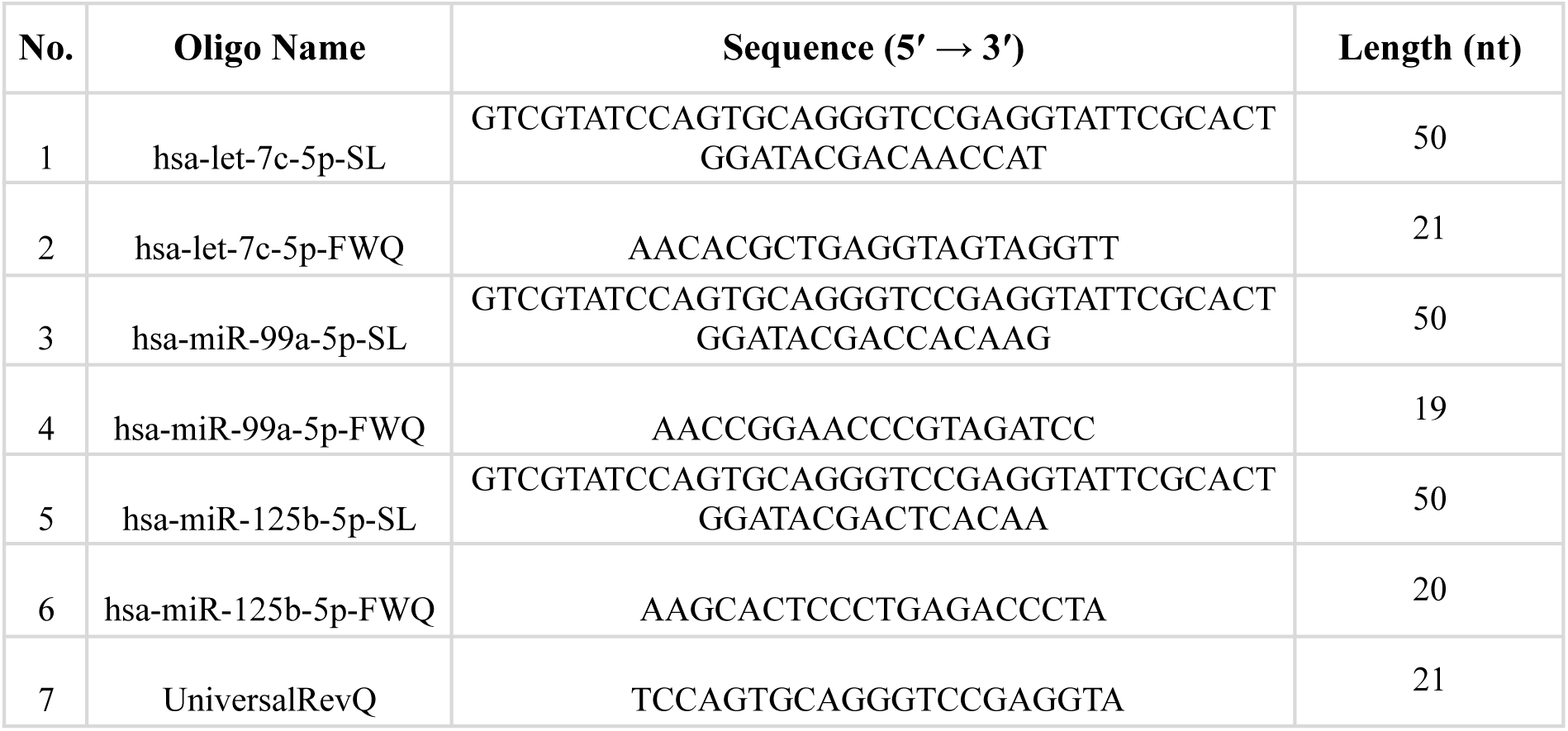
Primer Sequences Used for miRNA Reverse Transcription and qRT-PCR. (**Note:** All primers were desalted and synthesized without modifications. SL-Stem-loop primer; FWQ-Forward primer for qRT-PCR; UniversalRevQ-Universal reverse primer; hsa-Homo sapiens; nt-Nucleotides.)

### 2.4. Statistical analysis

#### Data Preprocessing

During the preprocessing phase, we identified and handled NULL values by replacing them with the median value in the qRT-PCR derived urinary exosomal miRNA expression data. All these operations were performed on the Ct values to ensure data consistency and readiness for analysis.

#### Statistical Analysis

We utilized the Mann-Whitney U test, a nonparametric approach, to evaluate the hypothesis. This test is beneficial since it does not rely on a normal distribution, making it versatile and effective for assessing differences between two independent groups. We used the ROC (Receiver Operating Characteristic) curve to assess the prediction potential of different characteristics in identifying HC and BCa classes. The ROC curve is a graphical tool for determining how well a binary model works at various threshold levels. The model’s performance at different threshold settings is plotted using two critical metrics: True Positive Rate (TPR) and False Positive Rate (FPR). The AUC (Area Under the Curve), which depicts the area below the ROC curve, is used to assess the model’s overall ability to correctly identify and discriminate between HC and BCa.

### 2.5. ML-Model Development

In this study, the Logistic Regression ML model has been employed as a baseline classifier for classifying BCa from HC, and also differentiating HG and LG cancer patients in BCa using demographic and qRT-PCR dataset to capture hidden and complex patterns. The model does not utilise any hyperparameter tuning for the classification task which further ensures the reproducibility of the proposed model. We have utilised the default hyperparameter values from scikit-learn library. To assess the models predictive performance we have utilized some performance matrix such as accuracy: Accuracy measures the overall correctness of a model. It is the fraction of all predictions (both positive and negative) that the model got right; precision: Precision focuses on the quality of positive predictions. It tells us, out of all the samples the model labeled as positive, how many were actually positive. High precision means fewer false alarms; recall: Recall focuses on the completeness of positive detection. It tells us, out of all the actual positive samples, how many the model successfully identified; F1 score: It is the harmonic mean of precision and recall. It balances the two, giving a single measure that is useful when both false positives and false negatives are important. Detailed steps of the model development process have been described in Figure 1B.

### 2.6. Bioinformatics Analysis

The target genes of 3 miRNAs were predicted using mirTarbase and Mirbase database. By removing duplicate entries all target genes list were merged, named as target gene panel (TGP) of miRNAs. Moreover, a panel of Bladder Cancer-specific genes (BCSG) panel were prepared using GeneCards, KEGG and STRING databases. The BCSG and TGP panels were crossed to identify common target genes (CTG) with Venny 2.0 (https://bioinfogp.cnb.csic.es/tools/venny/index2.0.2.html). These CTG represent the list of genes that have crucial roles in BCa molecular regulation and are regulated by let-7c cluster miRNAs. These were further explored for enrichment analysis, protein–protein interaction network construction, and pathway analysis to understand their roles in disease mechanisms. ShinyGO software was used for functional analysis of CTG panel. Cytoscape software was used for visualization and further analysis, allowing for hub genes extraction based on their crucial roles in the network.

## 3. Results

To evaluate the diagnostic potential of exosome-derived urinary miRNAs from the let-7c cluster in BCa, we analyzed their expression profiles using qRT-PCR in a well-characterized cohort. The study cohort consisted of BCa patients (n = 50) irrespective of tumor grade, age and sex-matched HC individuals (n = 16). The BCa cohort consisted of both HG (N=36) and LG (N=14) groups. Additionally, the BCa cohort was categorized by tumor stage into T1 (n = 32), T2 (n = 12) and Ta (n = 6). Their clinical and pathological features are summarized in **Table 2**.

**Table 2:** Clinical and general characteristics of BCa patients and HC.

| Biological Sample | *Urine* |  |
| --- | --- | --- |
| Variable | BCa (n=50) | HC (n=16) |
| Age (years, mean ± SD) | 59.1 ± 14.3 | 46.1 ± 15.0 |
| Gender | Male: 40 (80%)<br>Female: 10 (20%) | Male: 14 (87.5%)<br>Female: 2 (12.5%) |
| Stage | Ta: 6 (12%)<br>T1: 32 (64%)<br>T2: 12 (24%)<br>Carcinoma in situ: 0 (0%) | — |
| Grade | HG: 36 (72%)<br>LG: 14 (28%) | — |
| Tumor size | Mean: 4.41 ± 2.01<br>>3 cm: 34 (68%)<br>≤3 cm: 16 (32%) | — |
| Number of tumors | Single: 37 (74%)<br>Multiple: 13 (26%) | — |
| VI-RADS group | Low (1–3): 20 (40%)<br>High (4–5): 30 (60%) | — |
| Tobacco addiction | Yes: 31 (62%)<br>No: 19 (38%) | 0 (0%) |
| Alcohol intake | Yes: 12 (24%)<br>No: 38 (76%) | 0 (0%) |
| Diabetes mellitus | Yes: 10 (20%)<br>No: 40 (80%) | 0 (0%) |
| Hypertension | Yes: 8 (16%)<br>No: 42 (84%) | 0 (0%) |

### 3.1. Let7-c cluster gene expression pattern and correlation in BCa vs. HC

As shown in Figure 2, BCa patients exhibited a trend toward differential regulation of urinary exosomal let-7c-5p, miR-99a-5p and miR-125b-5p compared to HC. let-7c-5p expression was downregulated in BCa patients (P = 0.07621), miR-99a-5p and miR-125b-5p were also downregulated in BCa patients (P = 0.01327 and P = 0.08667, respectively). These results indicate that members of the let-7c cluster show a consistent trend of reduced expression in BCa relative to HC.

**Figure 2:**
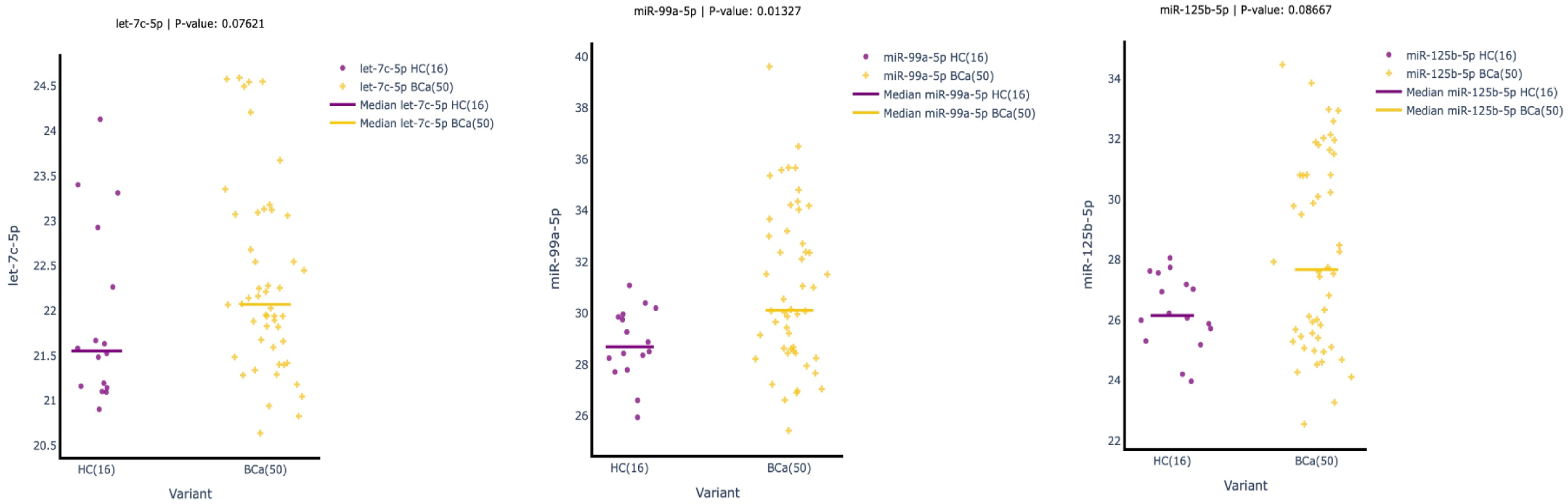
Expression levels of Let-7c cluster in urinary exosomes from HC (n=16) and BCa patients (n=50). Each dot represents an individual sample and horizontal lines indicate median values for each group.

Urinary exosomes from both cohorts were analyzed for the expression of let7-c cluster genes or miRNA using qRT-PCR. The heatmap was created using absolute Ct values obtained that show correlation between two cohorts (Figure 3). Pearson correlation analysis was used to assess relationships among urinary let-7c-5p, miR-99a-5p and miR-125b-5p in HC (Variant 0) and BCa patients (Variant 1). In HC, miR-99a-5p and miR-125b-5p showed a strong positive correlation (r = 0.73), suggesting possible co-regulation. In contrast, let-7c-5p displayed minimal correlation with either miR-99a-5p or miR-125b-5p, consistent with an independent expression pattern. Among cancer patients, correlations were weaker overall: miR-99a-5p and miR-125b-5p retained a moderate positive association (r = 0.54), while let-7c-5p shifted to weakly positive correlations with both miR-99a-5p (r = 0.37) and miR-125b-5p (r = 0.22). This attenuation of correlation in disease samples may reflect disruption of normal regulatory interactions within the let-7c cluster.

**Figure 3:**
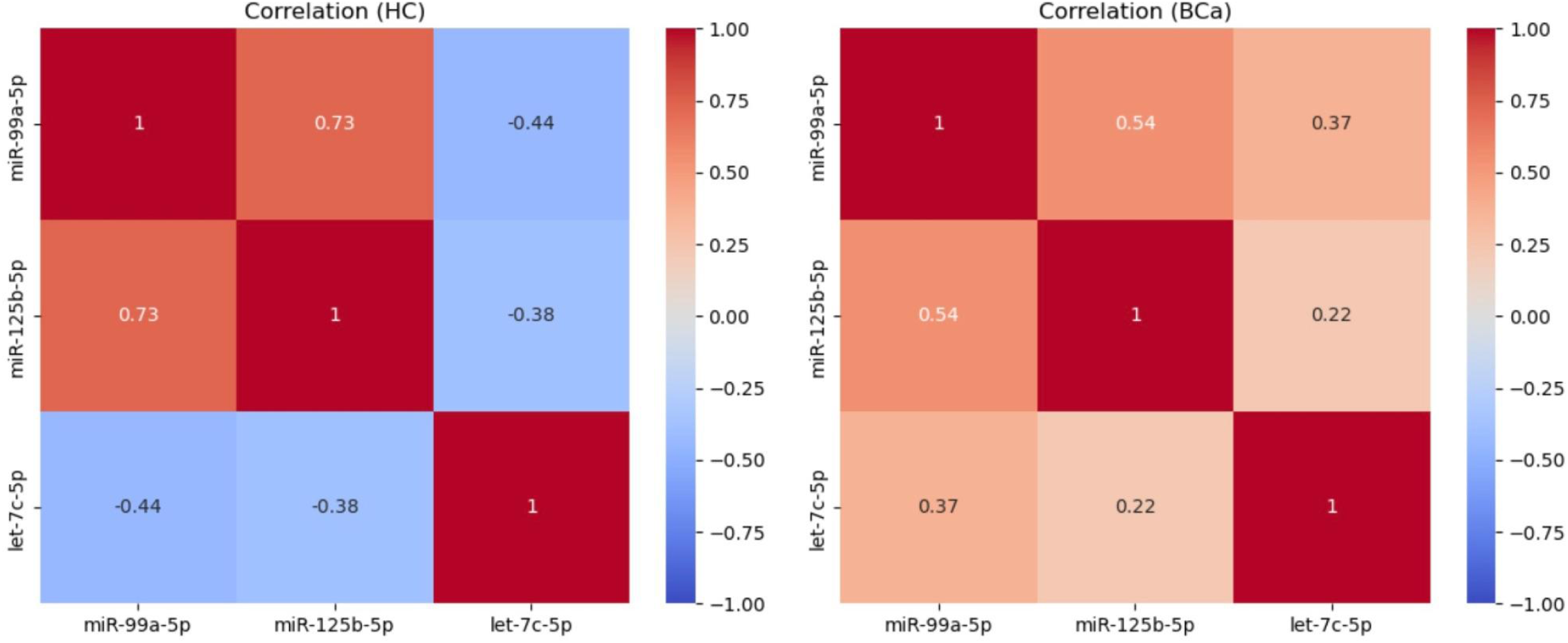
Correlation heatmaps of urinary miRNAs (let-7c-5p, miR-99a-5p, miR-125b-5p) in ‘HC’ and ‘BCa’ groups. Stronger positive correlations are seen in ‘HCs’, while ‘BCa’ shows weaker, more uniform positive associations, suggesting context-dependent miRNA co-regulation.

To demonstrate the discriminative power of individual urinary miRNAs in distinguishing BCa patients from HC, AU-ROC analysis was performed (Figure 4A). Among these three features, miR-99a-5p exhibited the highest diagnostic accuracy with an AU-ROC of 0.71, followed by let-7c-5p (AU-ROC = 0.65) and miR-125b-5p (AU-ROC = 0.64). In addition, the diagnostic potential of the ratio of these miRNAs (such as miR-99a-5p / let-7c-5p: AU-ROC=0.32; miR-125b-5p / let-7c-5p: AU-ROC=0.64; and miR-99a-5p / mir-125b-5p: AU-ROC=0.58;) were also evaluated using AU-ROC to explore combinatorial diagnostic strength.

**Figure 4.**
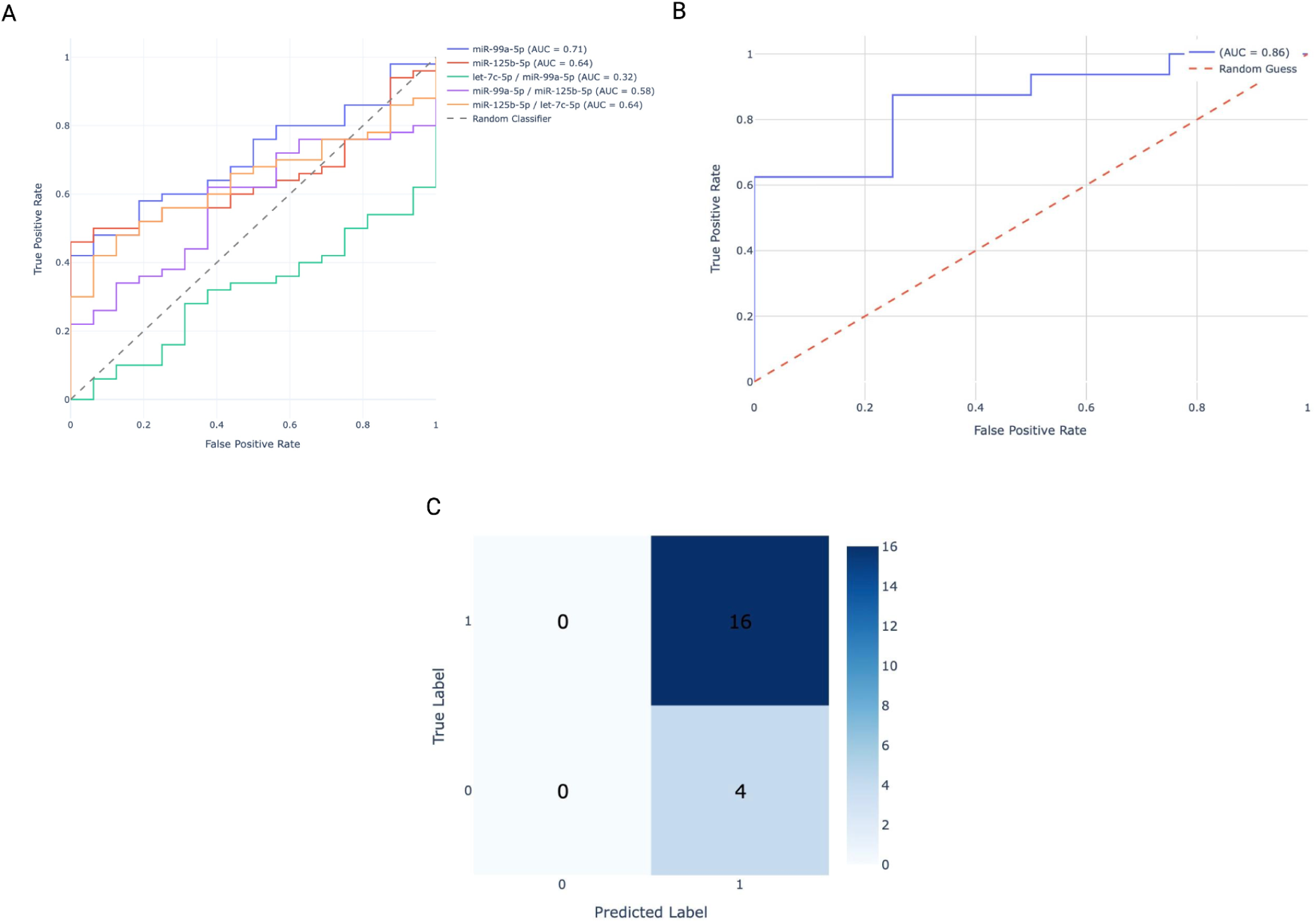
**(A)** ROC graph of individual features, **(B)** ROC graph of Logistic Regression model (HC vs BCa classification), **(C)** confusion matrix for the HC vs BCa classification.

To explore the scope of ML in this analysis, we employed logistic regression - a ML based approach. The cohort was divided into training (70%) and testing (30%) multiple features including individual miRNA: let-7c-5p, miR-99a-5p, miR-125b-5p; and ratio based features: miR-99a-5p_miR-125b-5p, miR-125b-5p_let-7c-5p and let-7c-5p_miR-99a-5p were used. The model achieved an AU-ROC of 0.86, indicating improved discriminative power as compared to individual features. The curve **Figure (4B)** shows a steep ascent towards the top-left quadrant, reflecting a high true positive rate (sensitivity) with minimal false positive classifications. The model achieved an accuracy of 80%, indicating it correctly classified 80% of the samples. With a precision (explain precision here) of 80% and a recall of 100%, it identified all true positive cases without missing any of it Figure 4(C). The F1-score of 0.889 reflects a good balance between precision and recall. The confusion matrix shows no false negatives but includes some false positives.

### 3.2. Comparison Based on Tumor Grades (HG vs. LG)

Using the same approach, we applied a logistic regression model to distinguish HG, n=36 from LG, n=14 cases with a 70/30 training–testing split. The model achieved an accuracy of 53.3% and an AU-ROC of 0.63, indicating only modest discriminatory power, close to random classification. To investigate why the model underperformed, especially in the context of tumor grades, we next evaluated whether miRNA expression patterns were influenced by clinical and lifestyle variables that could contribute additional discriminatory value. **Table 3** reports the p-value using two different tests (t-test and Chi-square test) which are between the tumour grade and all the clinical and miRNA features.

**Table 3:**
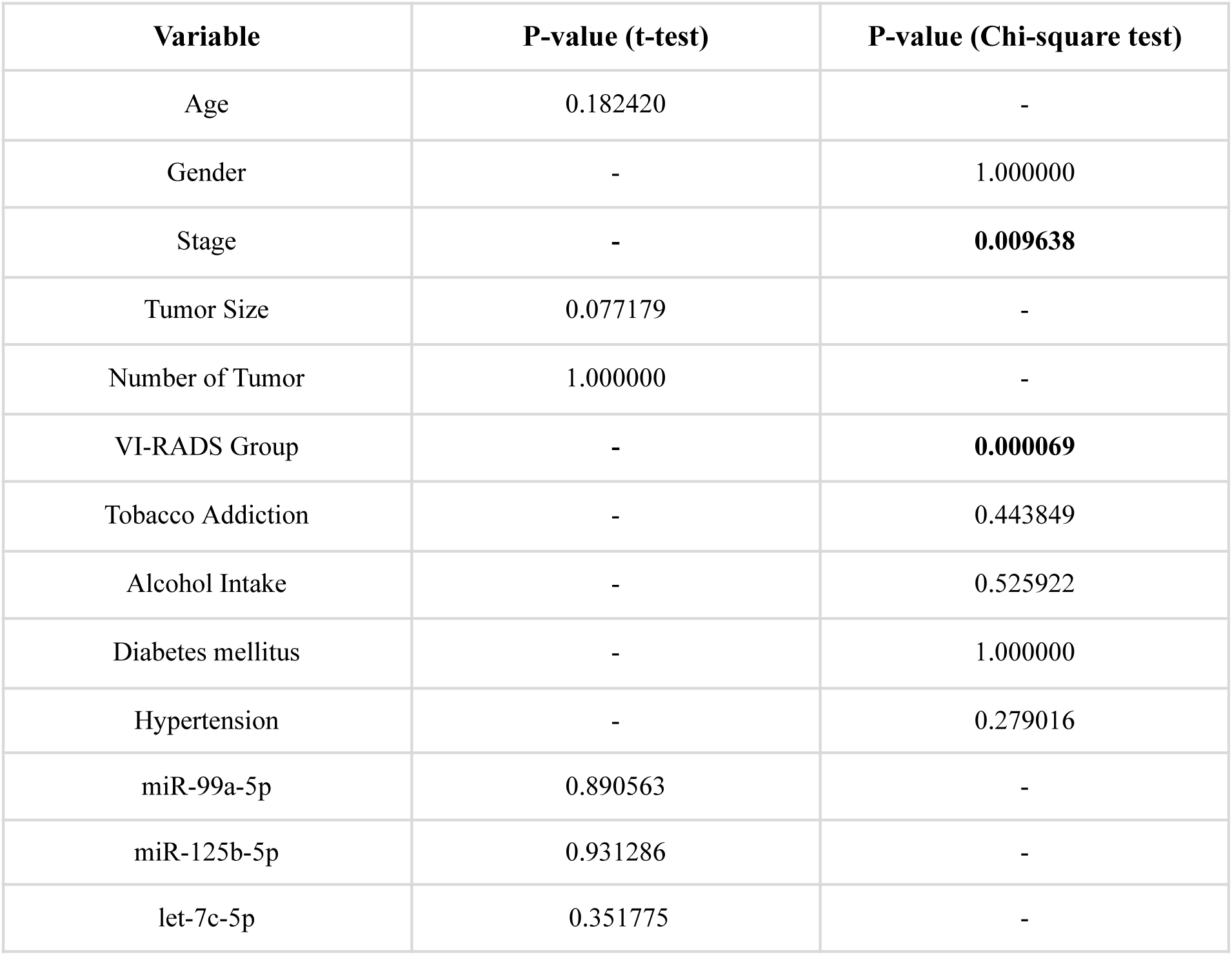
Hypothesis test result between Grade of cancer and all clinical variables and biomarkers.

| Variable | P-value (t-test) | P-value (Chi-square test) |
| --- | --- | --- |
| Age | 0.182420 | - |
| Gender | - | 1.000000 |
| Stage | - | **0.009638** |
| Tumor Size | 0.077179 | - |
| Number of Tumor | 1.000000 | - |
| VI-RADS Group | - | **0.000069** |
| Tobacco Addiction | - | 0.443849 |
| Alcohol Intake | - | 0.525922 |
| Diabetes mellitus | - | 1.000000 |
| Hypertension | - | 0.279016 |
| miR-99a-5p | 0.890563 | - |
| miR-125b-5p | 0.931286 | - |
| let-7c-5p | 0.351775 | - |

In the ML model, when miRNA data was combined with other clinical features showed the possibility of a potential improvement in the overall performance while distinguishing tumour grade. AU-ROC (Figure 5A) of 0.61 in classifying HG versus LG BCa cases using a combined feature set consisting of urinary miRNA expression (let-7c-5p, miR-99a-5p and miR-125b-5p) alongside key clinical and other variables. ML model was improved by 20% demonstrating accuracy of 73.33% presented in the confusion matrix (Figure 5B). However, the model achieved an AU-ROC of 0.61, indicating only modest discriminatory ability between HG and LG subgroups. While the AU-ROC curve slightly deviates from the diagonal line (0.50), the relatively flat slope and lower sensitivity values suggest that the current combination of features lack sufficient predictive power for reliable tumor grade differentiation.

**Figure 5:**
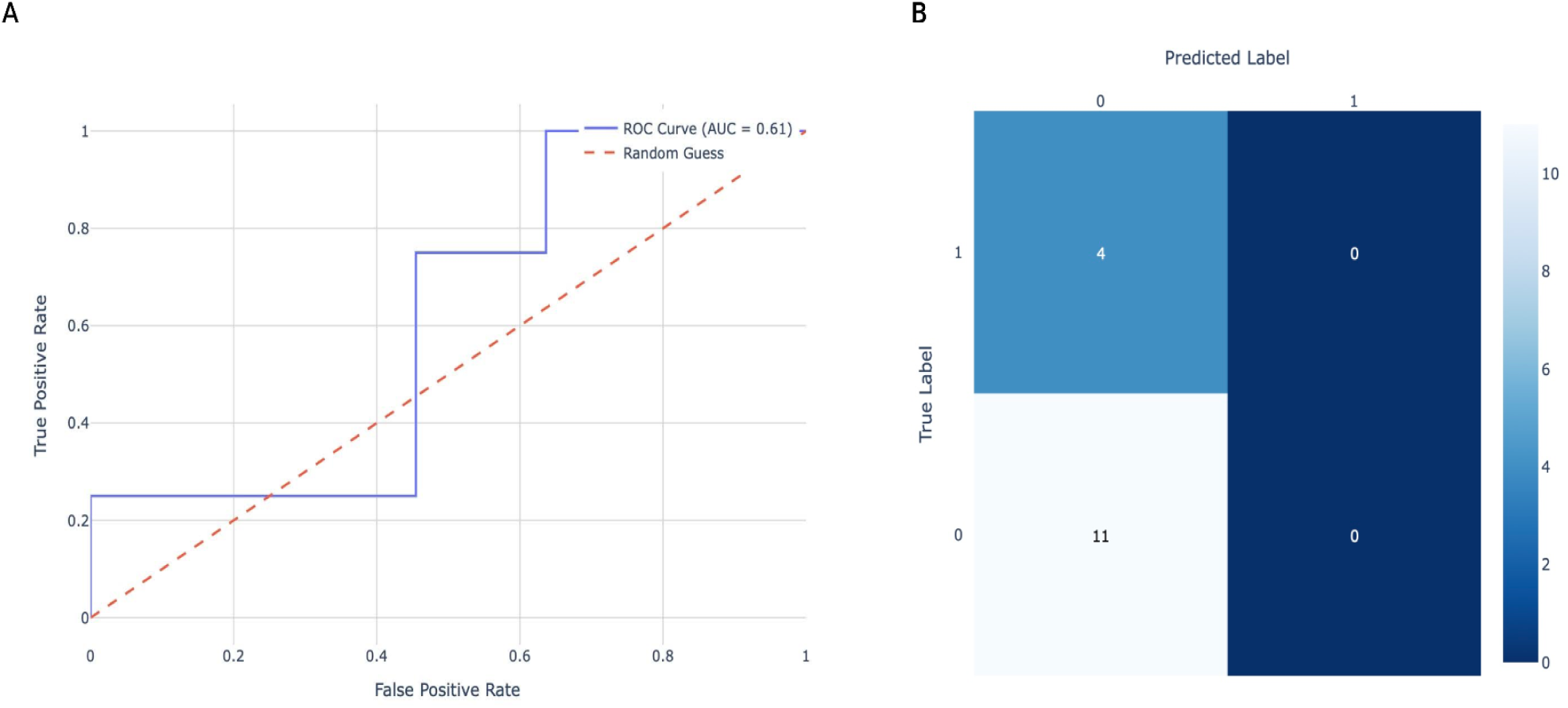
Results of Logistic Regression Model including molecular and clinical features as well to differentiate between HG and LG Cancer Patients. **(A)** ROC **(B)** Confusion Matrix

### 3.3. Let7c cluster expression pattern in Tumor Stages (Ta, T1, T2)

Let-7c-5p showed little stage-related variation and poor classification (AU-ROC 0.32–0.36) (Figure 6A**,B**), while miR-125b-5p also demonstrated limited stage-specific value (AU-ROC 0.43–0.64). In contrast, miR-99a-5p displayed clearer differences, especially at Ta and achieved stronger discriminatory potential (AU-ROC 0.79). These findings position miR-99a-5p as the most promising of the three markers for stage evaluation, though validation in larger cohorts will be required.

**Figure 6:**
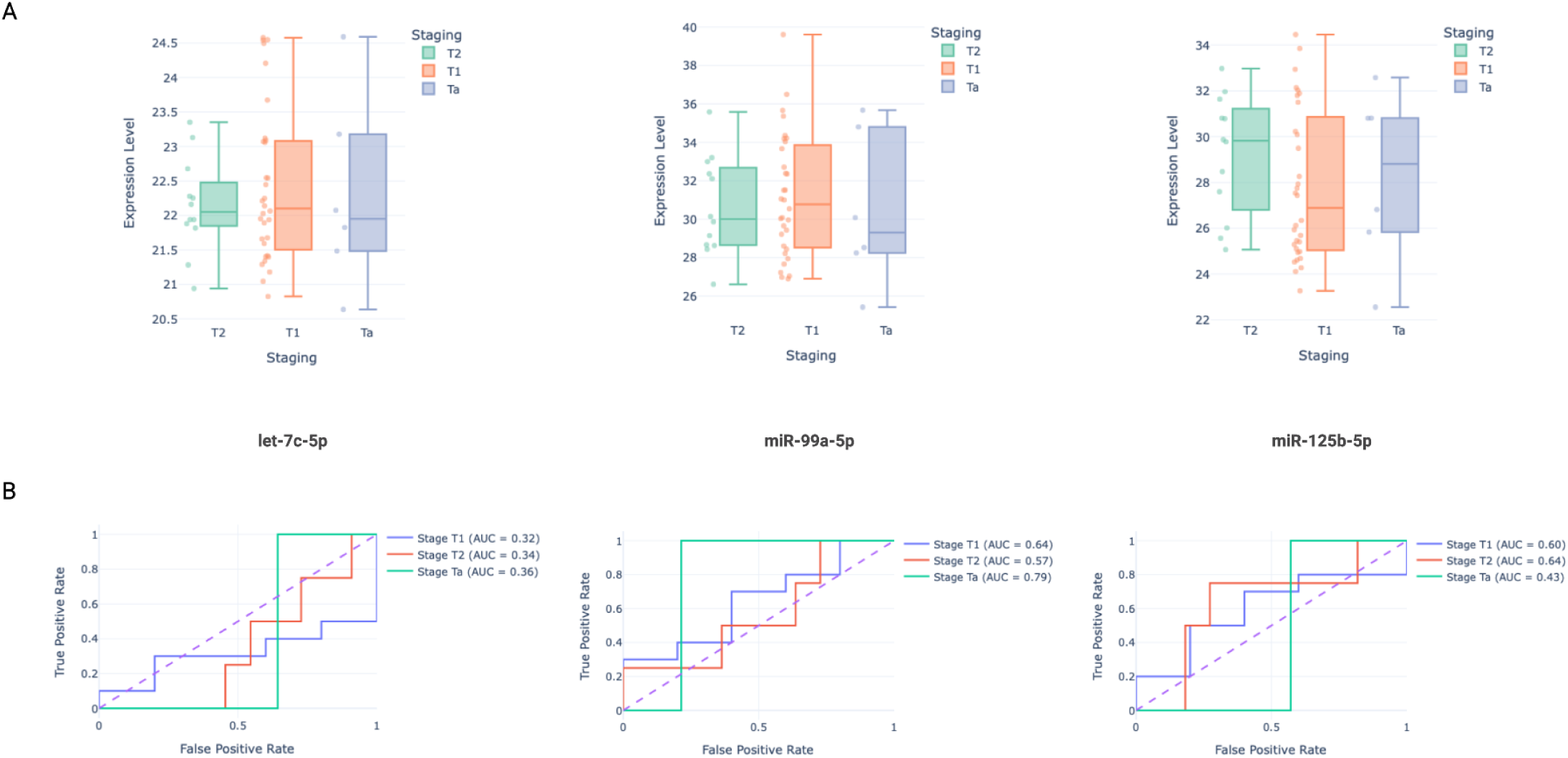
Boxplot and ROC of stage-specific classification: **(A)** Boxplot **(B)** ROC.

### 3.4. Validation of results in Bioinformatics Analysis

We identified target genes of all 3 miRNAs (Let-7c-5p, miR-99a-5p and miR-125b-5P) and predicted 3072 target genes (TGP) using mirTarbase^46^ and Mirbase database^47^. The segregation of target genes of miRNAs and BCSG panel are shown in Figure 7A. Further, TGP and BCSG panels were crossed and 109 CTG genes that were common in both panels were selected for further analysis. The PPI network was created using CTG genes in the STRING database. To identify interaction between the target genes of the miRNAs highest confidence threshold (≥0.9) and disconnected nodes were excluded to refine PPI network. Using Cytohubba tools^48^, 10 highly scored genes that showed the shortest path according to degree were identified, Figure 7B. We identified 10 hub gene: ACTB, BCL2, CCND1, CDKN2A, ERBB2(also known as HER2), IL6, MYC, STAT3, TNF, TP53, all showed association and play critical role in BCa pathogenesis. TP53, MYC and ACTB are top central hub genes (red nodes), while genes like STAT3, BCL2 and CCND1 show moderate connectivity (orange nodes), CDKN2A, ERBB2, IL6 and TNF were still involved with fewer direct interactions (yellow nodes). The hub genes were reanalyzed in the STRING database using a high-confidence threshold (≥ 0.9) to ensure accurate and biologically relevant protein–protein interactions, to avoid false positive and weak interaction. The disconnected nodes were removed to identify core genes central to BCa progression Figure 7C. The chord diagram analysis (Figure 8) shows the regulatory interactions between dysregulated miRNAs (let-7c-5p, miR-99a-5p and miR-125b-5p) along with their target genes.Some miRNA regulates multiple target genes and several genes are controlled by more than one miRNA, which signify regulatory links within the network. The shared regulation shows a cooperative approach, where multiple miRNAs work together to manage gene expression more effectively. Cross-connections add redundancy to the system, which helps maintain stable gene expression even when individual regulatory pathways are disrupted. This network shows the complexity of post-transcriptional regulation and identifies key genes that play important roles in biological processes and BCa.

**Figure 7:**
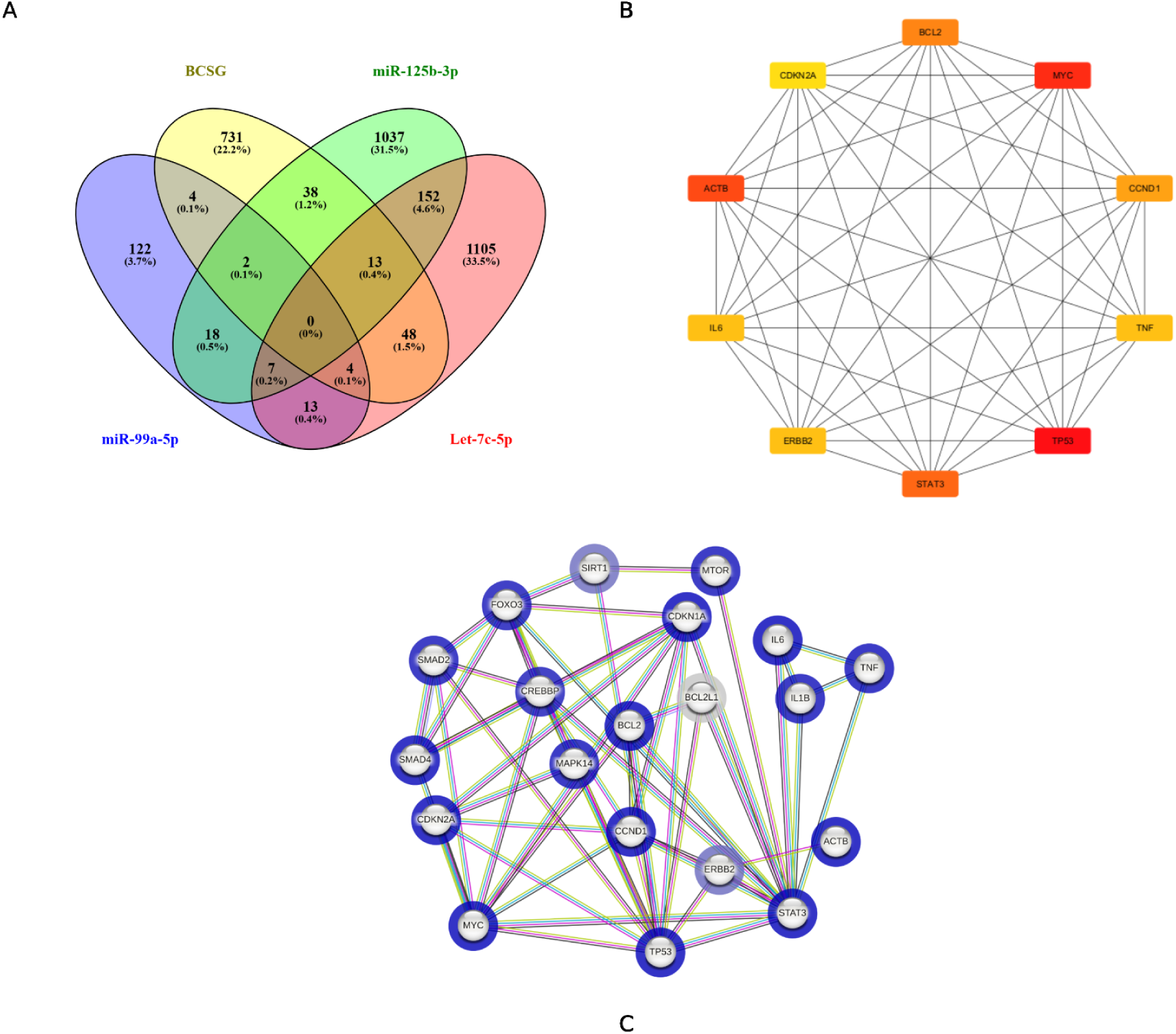
**(A)** Venn diagram showing miRNA and their target genes; common genes between the BCGS panel and miRNA gene targets were highlighted, **(B)** Top 10 genes with the highest degree scores were selected using CytoHubba tools, **(C)** miRNA–target genes and hub genes were reanalyzed using the STRING database,shown as a combined visualization.

**Figure 8:**
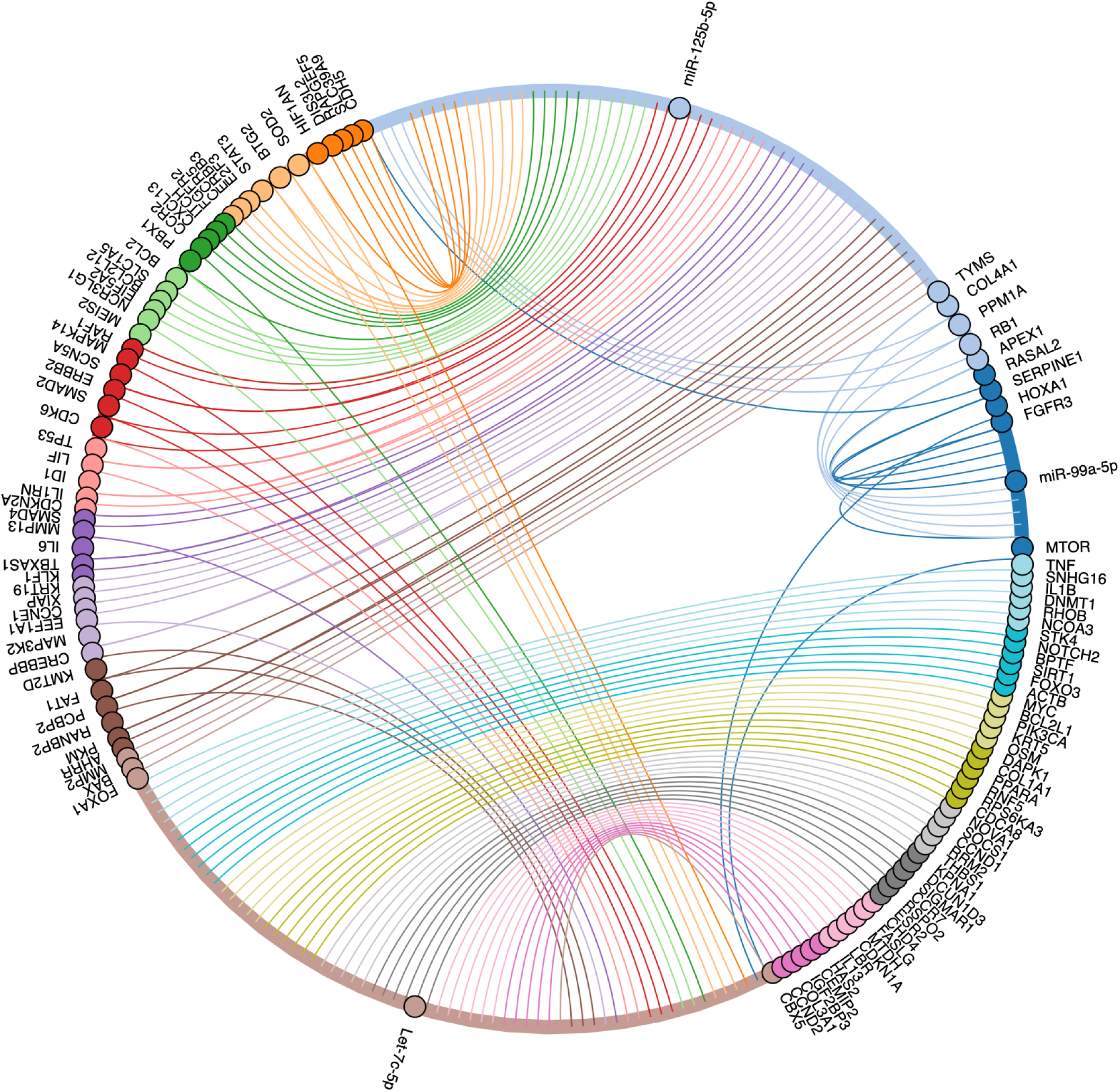
Chord diagram analysis of let-7c cluster and their gene target involved in BCa.

In biological process analysis Figure 9 **(A)**, the majority of genes are enriched in the response to endogenous stimuli and oxygen-containing compounds, apoptotic process regulation, programmed cell death regulation, and cell population proliferation regulation. Fibroblast growth factor binding, NF-kappaB binding, growth factor binding, and RNA polymerase II-specific DNA-binding transcription factor binding were all involved in the molecular function Figure 9 **(B)**. Cyclin-dependent protein kinase holoenzyme complex, serine/threonine protein kinase complex, protein kinase complex, nuclear membrane and envelope were the most abundant cellular components Figure 9 **(C)**. Using the KEGG database Figure 9 **(D)**, significant signaling pathways were identified, indicating that candidate genes were mostly involved in P53 signaling, cancer pathways, non-small cell BCa,lung cancer, pancreatic cancer and endocrine resistance.

**Figure 9:**
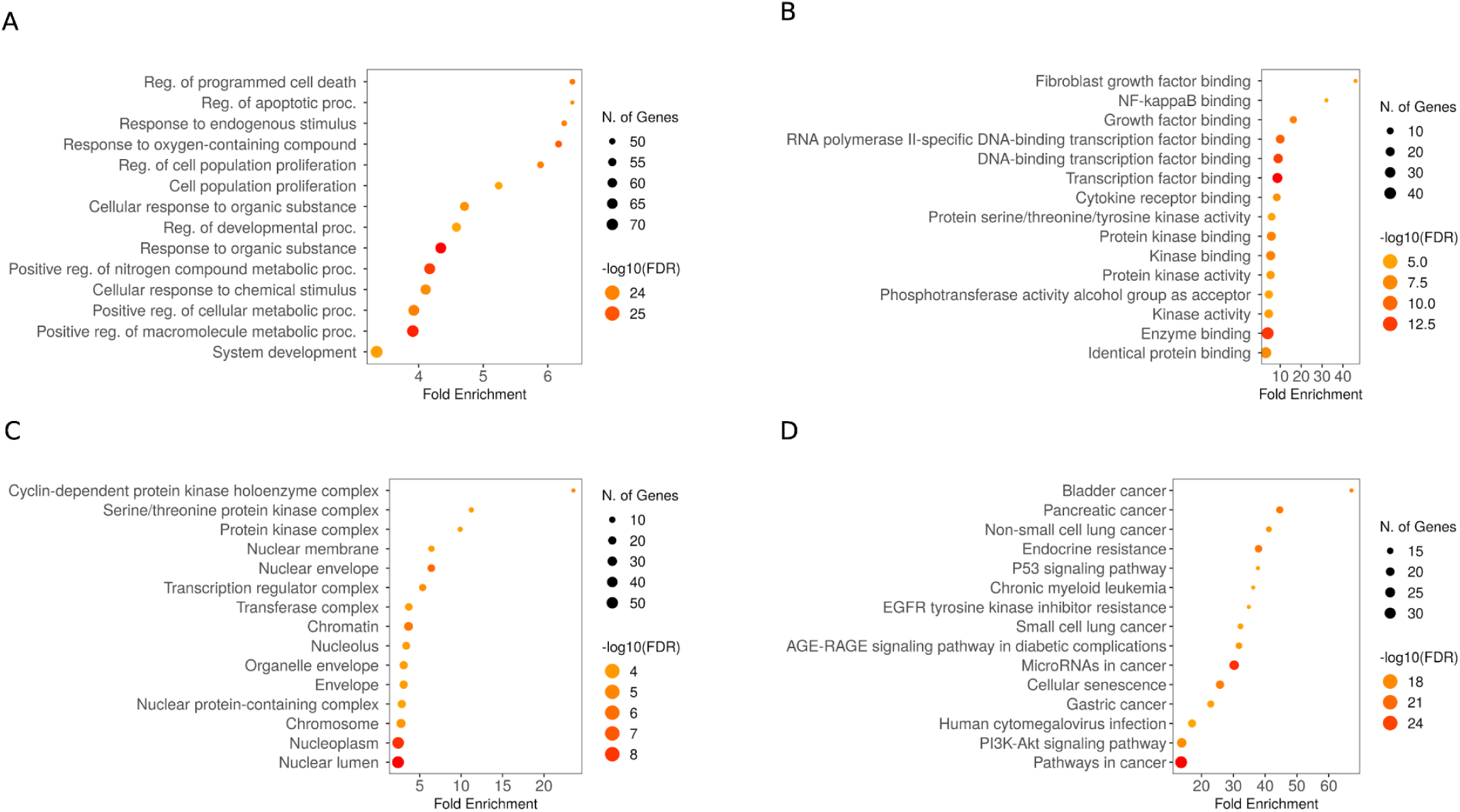
**(A)** Biological processes: illustrates metabolic pathways regulation essential for cellular processes and tumorigenesis, **(B)** Molecular function: shows crucial roles in transcription factor binding and kinase activity, **(C)** Cellular components: show significant roles in regulating gene expression and modulating cellular signaling, **(D)** KEGG pathway enrichment analysis: shows key cancer-related pathways emphasizing fold enrichment, significance, and gene involvement.

## 4. Discussion

This study demonstrates that urinary exosomal miRNAs from the let-7c cluster, particularly miR-99a-5p, can distinguish BCa patients from HC with high accuracy when analyzed using supervised ML. This is clinically relevant given the persistent challenge of differentiating BCa from benign urological conditions such as cystitis or urinary tract infection, which often present with overlapping symptoms like hematuria^49^. Current gold-standard diagnostic tools cystoscopy and urinary cytology remain invasive or insensitive, underscoring the need for reliable, non-invasive biomarkers. By showing that qRT-PCR derived miRNA data can be enhanced with ML to detect subtle expression changes, our findings contribute to efforts to translate molecular signatures into practical diagnostic tools.

### Comparison with current diagnostic modalities

In routine clinical evaluation, patients with hematuria are assessed with urinalysis, imaging, cytology and cystoscopy^50^. Imaging modalities such as ultrasound and CT urography can identify larger or infiltrative tumors but lack resolution for flat or early-stage lesions and MRI is mainly used for staging rather than initial diagnosis^51^. Urine cytology, while highly specific (∼78–86%), has poor sensitivity for LG tumors (∼16%) and only moderate sensitivity for HG disease (∼84%)^52^. Several FDA approved urine assays, including UroVysion FISH, ImmunoCyt/uCyt+ (Immunocytology test), NMP22 and BTA tests, have attempted to fill this gap but are limited by variable sensitivity and frequent false positives in benign conditions^53^. Our results suggest that exosomal miRNAs, when analyzed with ML, may overcome some of these limitations by offering biological specificity, resistance to degradation and the ability to capture subtle patterns not detectable with conventional approaches.

### Expression patterns and diagnostic performance

We observed consistent downregulation of the let-7c-5p/miR-99a-5p/miR-125b-5p in BCa patients compared with HC, with miR-99a-5p emerging as the most robust diagnostic candidate across grade and stage analyses. Let-7c-5p and miR-125b-5p also showed downregulation, though less prominently, but may hold value in ratio-based features that reduce sample variability and capture biologically relevant interactions. Correlation analysis supported this view: in HC, miR-99a-5p and miR-125b-5p were strongly co-expressed, whereas let-7c-5p was largely independent, suggesting distinct regulatory mechanisms. These coordinated patterns were attenuated in cancer samples, consistent with disruption of normal regulatory interactions. Such findings highlight why ratio-based and multimodal features are advantageous in diagnostic modeling. In addition, the altered correlation patterns suggest a loss of coordinated regulation in the cancer state, highlighting potential changes in molecular interactions between healthy and malignant conditions.

### Relation to prior studies

Our results both align with and diverge from prior studies. Spagnuolo et al. described significant upregulation of urinary let-7c in HG NMIBC, particularly T1 tumors, with strong stage-discrimination performance (AU-ROC= 0.80)^32^. While their study focused on prognostic risk stratification following BCG treatment in HG NMIBC, we observed downregulation of the Let-7c cluster in BCa across a broader clinical spectrum. Although stage-specific discrimination in our cohort was modest, these discrepancies may reflect differences in study design, cohort size or clinical context (prognostic vs diagnostic focus). Importantly, by integrating expression ratios and ML, we extended the clinical applicability of the cluster beyond prognostic stratification to diagnostic classification. Consistent with this, Zhang et al. demonstrated that urinary miR-99a and miR-125b are significantly downregulated in BCa, achieving AU-ROC values of 0.800 and 0.813, respectively, with a combined AU-ROC of 0.876, underscoring their diagnostic utility as non-invasive biomarkers^44^. In our study, miR-99a-5p emerged as the most consistent biomarker associated with disease presence, while let-7c-5p contributed more as a relatively low-expressing but stable biomarker within ratio-based features. In contrast to its role as a primary classifier in previous prognostic approaches, let-7c-5p in our study serves a more refined purpose as a stable reference point within ratio features. This diverse functional role of let-7c-5p aligns with our earlier work, where we introduced our in-house algorithm for reference miRNA selection and identified let-7c-5p as a highly stable urinary exosomal miRNA across multiple urological conditions^54,55^.

The downregulation of the let-7c cluster has been frequently observed in various cancer types, where its loss is commonly associated with tumor-suppressive functions and poor clinical outcomes^28,56–58^. Our findings are consistent with previous research, revealing that let-7c-5p is downregulated in BCa. However, its diagnostic ability was weaker to that of miR-99a-5p and miR-125b-5p, which were more effective at distinguishing BCa patients from HC. Previous studies have similarly reported reduced urinary expression of let-7c-5p in BCa, reinforcing its role as a tumor suppressor^59,60^. In contrast, Spagnuolo et al.’s findings of let-7c-5p upregulation in HG NMIBC further emphasize the context-dependent variability of its expression patterns^32^. miR-99a-5p emerged as the most reliable diagnostic marker in our study, a finding supported by multiple studies demonstrating its consistent downregulation in BCa tissues and urine^44,61–63^. Additionally, miR-125b-5p has been consistently reported as downregulated in BCa, strengthening its tumor-suppressive role^44,64–67^.

### ML integration

Logistic regression was chosen for its interpretability, robustness with small datasets and ability to generate probability estimates relevant to clinical decision-making. Bitin et al. employed a multi-modal approach, combining miRNA data with clinical variables and achieving an AU-ROC of 0.85. While their study utilized a broader set of miRNAs and clinical data, we demonstrated similar diagnostic accuracy (AU-ROC = 0.86, sensitivity 100%, accuracy 80%) and clear separation between BCa patients and HC with moderate grade classification through a more focused approach using qRT-PCR derived miRNA data and logistic regression. The inclusion of molecular and clinical variables improved diagnostic precision, though this may have been influenced by sample size and class imbalance. The individual AU-ROC values of single miRNAs for stage discrimination were very low, further emphasizing the need for integrated models. These results should therefore be considered proof-of-concept rather than clinically validated. Further optimization using larger, balanced cohorts and more advanced algorithms such as ensemble or deep learning approaches may enhance performance, particularly for grade and stage classification.

### Biological validation through bioinformatics

To ensure interpretability and biological relevance, we conducted integrative bioinformatics analyses. KEGG enrichment revealed associations of miR-99a-5p with PI3K–Akt, mTOR, and FGFR3 pathways, miR-125b-5p with p53, NF-κB, and HIF-1 signaling, and let-7c-5p with RAS/MAPK and cell cycle regulation. These pathways are central to BCa progression, underscoring the functional plausibility of our findings. PPI network analysis identified hub genes such as TP53, MYC, STAT3, EGFR, IGF1R and CCND1, reflecting the interplay of oncogenes and tumor suppressors in BCa biology. The coexistence of both stimulatory and inhibitory regulators illustrates why single miRNAs have limited discriminatory power, and why combined, ML-enabled signatures are needed to capture net biological effects. GO enrichment highlighted processes including DNA damage response, EMT, and immune modulation, further linking the cluster to key hallmarks of BCa. By directly regulating important oncogenic pathways like FGFR3 and mTOR, miR-99a-5p functions as a tumor suppressor biologically^68,69^, thereby enhancing its biological significance and translational potential as a credible non-invasive biomarker. Collectively, these results imply that while urinary levels of the let-7c cluster may differ depending on tumor grade, disease stage or methodological considerations, our study’s consistent downregulation, which is consistent with other reports, highlights the biological significance and diagnostic accuracy of this cluster in BCa.

### Limitations and future directions

This study has several limitations. The single-center cohort was modest in size, limiting statistical power for grade- and stage-specific analyses and reducing generalizability. Validation in larger, multi-center cohorts is required. The biomarker panel was restricted to three miRNAs and may not capture the full molecular complexity of BCa. Broader high-throughput profiling, coupled with functional validation, could refine biomarker panels and clarify mechanistic roles. Biological heterogeneity in urinary exosomes, which carry material from both malignant and non-malignant cells, also remains a challenge. Standardized protocols for exosome isolation, normalization, and qRT-PCR analysis will be essential for reproducibility across studies. From a modeling perspective, external validation is necessary to establish clinical robustness, and future work should test more advanced ML frameworks. Finally, our cross-sectional design precluded evaluation of prognostic utility; longitudinal studies are needed to determine whether these miRNAs can predict recurrence, progression, or treatment response. Integration with other modalities, such as mpMRI or VI-RADS, may further improve diagnostic precision and clinical adoption. At the same time, the biological heterogeneity of urinary exosomes carrying material from both malignant and non-malignant cells remains a challenge for reproducibility. Future studies should therefore focus on methods and tools to enrich tumor-derived exosomes. These results suggest that, when combined with ML techniques, urinary exosomal miRNA-based diagnostics offer a more precise and clinically applicable method for BCa detection and stratification. This approach could potentially reduce the need for invasive procedures while maintaining high sensitivity. Although this represents a significant advancement, the low AU-ROC emphasizes the need for further optimization. This could be achieved by adding more features to the model or adopting advanced model frameworks, such as ensemble methods or deep learning techniques, which could potentially enhance diagnostic accuracy.

## 5. Conclusion

This study provides proof of concept that urinary exosomal miRNAs, particularly miR-99a-5p, can contribute to BCa detection when analyzed within a ML framework. By integrating molecular and clinical features, our approach demonstrated reliable discrimination between BCa & HC and highlighted the value of ratio-based and multimodal features for diagnostic refinement. While current performance reflects the limitations of sample size and scope, these findings reinforce the promise of urinary exosomal miRNAs as non-invasive biomarkers and establish a foundation for larger validation studies. Future efforts should focus on expanding biomarker panels, standardizing analytical pipelines and combining molecular signatures with imaging and clinical parameters to advance toward clinically deployable diagnostic tools.

## 6. Contributions

*Research conceptualization and design*: Kumar L, Jain G and Kural S; *Supervision and Validation*: Kumar L and Jain G; *Project Administration*: Kumar L; *Collection and/or assembly of data*: Kural S, Pathak AK, Singh S and Kumar L; *miRNAs analysis workflow*: Kural S; *Model Engineering/Development*: Pathak AK and Gupta M; *Data analysis and interpretation*: Pathak AK, Kural S, Singh S and Kumar L; *Visualization*: Kural S, Pathak AK and Singh S; *Writing-Original Draft*: Kural S, Pathak AK, Singh S, Jain G and Kumar L; *Resources*: Yadav M, Kumar U, Singh Y, Trivedi S and Das P; *Writing-Review & Editing*: Kumar L, Jain G, Gautam V, Gupta M, Das P, Kural S, Pathak AK and Singh S; *Funding Acquisition*: Kumar L

## 9. Informed Consent

Every participant received comprehensive information about the study and their participation was completely voluntary. Written consent was obtained before any data was collected.

## 10. Data Availability Statement

We are dedicated to promoting reproducibility and openness in research. Anonymous datasets and supplementary information used in this study can be shared for research or verification purposes, but access to patient-related data is limited to protect privacy and meet ethical standards. Subject to institutional approval and suitable data-sharing agreements, interested researchers can get in touch with the corresponding author and data will be made available upon reasonable request.

## Acknowledgement

We would like to acknowledge the Centre for Genetic Disorders, Institute of Science, Banaras Hindu University (BHU) for providing essential laboratory facilities and research support throughout the course of this study. We are also grateful to the Department of Urology, Institute of Medical Sciences (IMS), BHU for their invaluable assistance in coordinating and facilitating sample collection. We also appreciate the support of the BioNEST, Central Discovery Centre, BHU for access to additional laboratory resources. We acknowledge the support of the Department of Computer Science, Institute of Science, BHU in developing and refining the ML models. We are sincerely thankful to all collaborators and the patients whose participation made this study possible.

## 11. Abbreviations

ACTB: Actin Beta
AI: Artificial Intelligence
AUC: Area under the curve
AU-ROC: Area Under the Receiver Operating Characteristic
BCa: Bladder Cancer
BCG: Bacillus Calmette-Guérin
BCSG: Bladder Cancer-Specific Genes
BCL2: B-cell Lymphoma 2
BTA: Bladder Tumor Antigen
CCND1: Cyclin D1
CDKN2A: Cyclin Dependent Kinase Inhibitor 2A
cDNA: complementary DNA
c-MYC: Cellular Myelocytomatosis Oncogene
CT: Computed Tomography
CTG: Common Target Genes
DMTF1: Cyclin D Binding Myb-Like Transcription Factor 1
EGFR: Epidermal Growth Factor Receptor
EMT: Epithelial–Mesenchymal Transition
ERBB2 / HER2: Erb-B2 Receptor Tyrosine Kinase 2 / Human Epidermal Growth Factor Receptor 2
EVs: Extracellular Vesicles
FDA: U.S. Food and Drug Administration
FGFR3: Fibroblast Growth Factor Receptor 3
FISH: Fluorescence In Situ Hybridization
FPR: False Positive Rate
GO: Gene Ontology
HIF-1: Hypoxia-Inducible Factor 1
HMGA2: High Mobility Group AT-Hook 2
H-Ras: Harvey Rat Sarcoma Viral Oncogene Homolog
IGF1R: Insulin-Like Growth Factor 1 Receptor
IL6: Interleukin 6
KEGG: Kyoto Encyclopedia of Genes and Genomes
LG: Low-Grade
MAPK: Mitogen-Activated Protein Kinase
miRNAs: microRNAs
Mirbase: microRNA Database
mirTarbase: microRNA–Target Interaction Database
MIBCa: Muscle Invasive Bladder Cancer
ML: Machine Learning
MRI: Magnetic Resonance Imaging
mTOR: Mammalian Target of Rapamycin
MYC: MYC Proto-Oncogene
NF-κB: Nuclear Factor kappa-light-chain-enhancer of activated B cells
NMP22: Nuclear Matrix Protein 22
NMIBCa: Non–Muscle Invasive Bladder Cancer
PDCD4: Programmed Cell Death 4
PI3K–Akt: Phosphoinositide 3-Kinase–Protein Kinase B signaling pathway
PPI: Protein–Protein Interaction
PTEN: Phosphatase and Tensin Homolog
qRT-PCR: Quantitative Real-Time PCR
RAS: Rat Sarcoma Viral Oncogene Homolog
RNA: Pol II RNA Polymerase II
ROC: Receiver Operating Characteristic
STAT3: Signal Transducer and Activator of Transcription 3
TNF: Tumor Necrosis Factor
TGP: Target Gene Panel
TP53: Tumor Protein P53
TPR: True Positive Rate
TURBT: Transurethral Resection of Bladder Tumor
UTIs: Urinary Tract Infections

